# Absence of Sac2/INPP5F enhances the phenotype of a Parkinson’s disease mutation of synaptojanin 1

**DOI:** 10.1101/2020.01.21.914317

**Authors:** Mian Cao, Daehun Park, Yumei Wu, Pietro De Camilli

**Author notes:** These authors contributed equally to this work (Mian Cao, Daehun Park). Program in Neuroscience and Behavioral Disorders, Duke-NUS Medical School 169857, Singapore.

## Abstract

Many genes whose mutations cause, or increase the risk of, Parkinson’s disease (PD) have been identified. An inactivating mutation (R258Q) in the Sac inositol phosphatase domain of synaptojanin 1 (SJ1/PARK20), a phosphoinositide phosphatase implicated in synaptic vesicle recycling, results in PD. The gene encoding Sac2/INPP5F, another Sac domain containing protein, was identified as a PD risk locus by GWAS. Knock-In mice carrying the SJ1 patient mutation (SJ1^RQ^KI) exhibit PD features, while Sac2 knockout mice (Sac2KO) do not have obvious neurological defects. We report a “synthetic” effect of the SJ1 mutation and the KO of Sac2 in mice. Most mice with both mutations died perinatally. The occasional survivors had stunted growth, died within 3 weeks, and showed abnormalities of striatal dopaminergic nerve terminals at an earlier stage than SJ1^RQ^KI mice. The abnormal accumulation of endocytic factors observed at synapses of cultured SJ1^RQ^KI neurons was more severe in double mutant. Our results suggest that SJ1 and Sac2 have partially overlapping functions and are consistent with a potential role of Sac2 as a PD risk gene.

## Introduction

Synaptojanin 1 (SJ1), also referred to as PARK20, is a phosphoinositide phosphatase concentrated at synapses implicated in endocytic membrane traffic. SJ1 dephosphorylates PI(4,5)P_2_ on endocytic membranes via the sequential action of two tandemly arranged inositol phosphatase modules: a central 5-phosphatase domain and an N-terminal Sac phosphatase domain (Fig 1A). The latter functions primarily as a 4-phosphatase although it can also dephosphorylate PI3P and PI(3,5)P_2_ in addition to PI4P *in vitro* (McPherson, Garcia et al., 1996).

**Figure 1.**
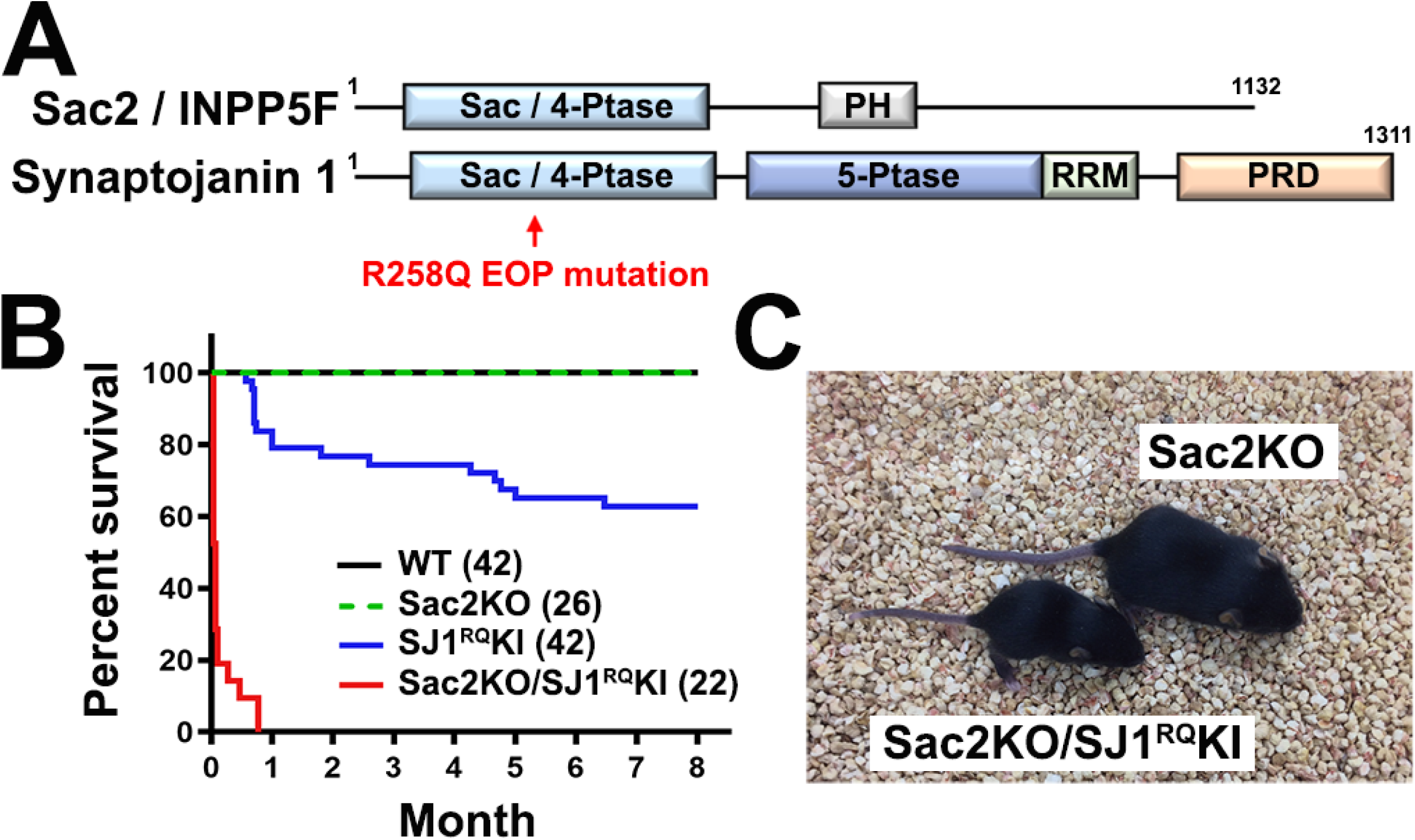
Genetic interaction between SJ1 and Sac2/INPP5F. A. Domain structures of SJ1 and Sac2. Sac/4-Ptase: 4-phosphatase domain; 5-Ptase: 5-phosphatase domain; RRM: RNA Recognition Motif; PRD: Proline-Rich Domain; PH: Pleckstrin Homology. B. Survival curves of mice with indicated genotypes. Numbers of animals are indicated in parenthesis. C. Littermate mice at postnatal day 15 (P15). The Sac2KO/SJ1^RQ^KI shown is one of the few survivors at this age.

Full loss-of-function mutations of the SJ1 gene result in early postnatal lethality in both mice and humans (Cremona, Di Paolo et al., 1999, Dyment, Smith et al., 2015, Hardies, Cai et al., 2016). In contrast, a homozygous missense R>Q mutation at a.a. position 258 within the Sac domain of SJ1 (a highly conserved a.a. position) is responsible for early-onset Parkinsonism (EOP) with epilepsy, leading to the naming of SJ1 as PARK20 (Krebs, Karkheiran et al., 2013, Olgiati, De Rosa et al., 2014, Quadri, Fang et al., 2013).

The main and well-established function of SJ1 is to participate in clathrin uncoating during the endocytic recycling of synaptic vesicles. SJ1 recruitment to endocytic pits by endophilin is required for dissociation of the PI(4,5)P_2_-dependent interaction of the clathrin adaptors with the membrane (Hayashi, Raimondi et al., 2008, Milosevic, Giovedi et al., 2011, Schuske, Richmond et al., 2003, Verstreken, Koh et al., 2003). Such action cooperates in uncoating with the disassembly of the clathrin lattice mediated by the triple AAA ATPase HSC70 and its synaptically enriched cofactor auxilin (Eisenberg & Greene, 2007, Fotin, Cheng et al., 2004). Interestingly, loss of function mutations in auxilin (PARK19) also result in EOP with epilepsy (Edvardson, Cinnamon et al., 2012, Koroglu, Baysal et al., 2013, Olgiati, Quadri et al., 2015). These observations support a link between dysfunction in clathrin mediated budding and EOP.

Previously, we have shown that the R258Q mutation abolishes the activity of the Sac phosphatase domain of SJ1 without affecting the activity of its 5-phosphatase domain (Krebs et al., 2013). We have also generated homozygous Knock-In (KI) mice (SJ1^RQ^KI mice) that carry the patient mutation and have neurological manifestations reminiscent of those of human patients (Cao, Wu et al., 2017a). Functional and microscopic analysis of cultured neurons and brain tissue of these mice demonstrated defects in synaptic vesicle recycling and dystrophic changes selectively in nerve terminals of dopaminergic neurons in the striatum (Cao et al., 2017a). Levels of auxilin (PARK19) and other endocytic factors were abnormally increased in the brains of such mice (Cao et al., 2017a). Additionally, levels of the PD gene parkin (PARK2) were strikingly elevated (Cao et al., 2017a) strengthening evidence for a functional partnership between endophilin, SJ1 and parkin revealed by previous studies (Cao, Milosevic et al., 2014, Matta, Van Kolen et al., 2012, Trempe, Chen et al., 2009).

Most interestingly, the intronic SNP rs117896735 located within the gene encoding Sac2/INPP5F, another protein containing a Sac domain which functions primarily as a PI4P 4-phosphatase, (Fig 1A) (Hsu, Hu et al., 2015, Nakatsu, Messa et al., 2015), was identified as a risk locus in PD by GWAS studies (Blauwendraat, Heilbron et al., 2019, Nalls, Pankratz et al., 2014). Sac2 has been implicated in the endocytic pathway via its interaction with Rab5 and, like SJ1, is preferentially expressed in the nervous system (Hsu et al., 2015, Nakatsu et al., 2015). These findings raise the hypothesis that the Sac domains of SJ1 and Sac2 may have some overlapping functions in the downregulation of a same phosphoinositide pool on membranes of the endocytic pathway at synapses. Here we used mouse genetics to explore this possibility. We report the occurrence of a strong synthetic effect both at the organismal level and at the synaptic level of the Sac2 null mutation and of the EOP mutation in the Sac domain of SJ1. Our results corroborate evidence for a link between impaired endocytic flow at synapses and EOP.

## Results

### Synthetic effect of SJ1 and Sac2 mutations on mice survival

Sac2KO mice do not have an obvious pathological neurological phenotype (Zou, Stagi et al., 2015). SJ1^RQ^KI mice are neurologically impaired and have seizures but about 60% of them can reach adulthood (Cao et al., 2017a). Mating of Sac2KO mice (Zhu, Trivedi et al., 2009) to SJ1^RQ^KI mice (Cao et al., 2017a) to generate animals homozygous for both mutations revealed a striking genetic interaction between Sac2 and SJ1. Sac2KO/SJ1^RQ^KI double mutant mice were born with mendelian ratio (Fig EV1A) but had a dramatically shorter lifespan than SJ1^RQ^KI mice (Fig 1B). Most of them died within 24 hours after birth, a few survived for a few days, but failed to thrive, and the remaining ones died before weaning within 3-weeks (Fig 1B and C). The Sac2 gene is adjacent to the Bag3 gene in both humans and mice, raising the possibility that a PD-relevant SNP found in the Sac2 gene (Nalls et al., 2014) may impact the expression of Bag3, a protein reported to impact synuclein clearance (Cao, Yang et al., 2017b). In the Sac2KO mice used for this study, however, brain levels of Bag3 were normal (Fig EV1B).

The striking synthetic effect of the lack of Sac2 with the inactivating mutation in the Sac domain of SJ1 raise the possibility that that Sac2 may have some overlapping function with SJ1. It was therefore of interest to determine whether a pool of Sac2 is present in axon terminals and is associated with endocytic structures. Available anti-Sac2 antibodies did not reveal a reliable immunoreactivity in either brain tissue or neuronal cultures in spite of the reported enrichment of Sac2 in brain (Hsu et al., 2015). Thus, we expressed GFP-Sac2 in WT cultured cortical neurons using either a lentivirus vector or calcium-phosphate transfection. GFP-Sac2 had a diffuse and broad distribution throughout neurons, including axon and axon terminals, which were visualized by anti-synaptophysin immunostain (Fig EV2) or mCherry-Rab3 overexpression (Fig 2A). Furthermore, when GFP-Sac2, which displays properties of a Rab5 effector (Hsu et al., 2015, Nakatsu et al., 2015), was co-expressed with mRFP-Rab5, a striking recruitment of GFP-Sac2 to mRFP-Rab5-positive structures was observed in axons (Fig 2B). Most likely the diffuse cytosolic localization in the absence of mRFP-Rab5 overexpression is due to saturation of endogenous Rab5. This cytosolic pool of GFP-Sac2 was not abolished by expression of mCherry-Rab3, These results make plausible an overlapping role of Sac2 and SJ1 in nerve terminals.

**Figure 2.**
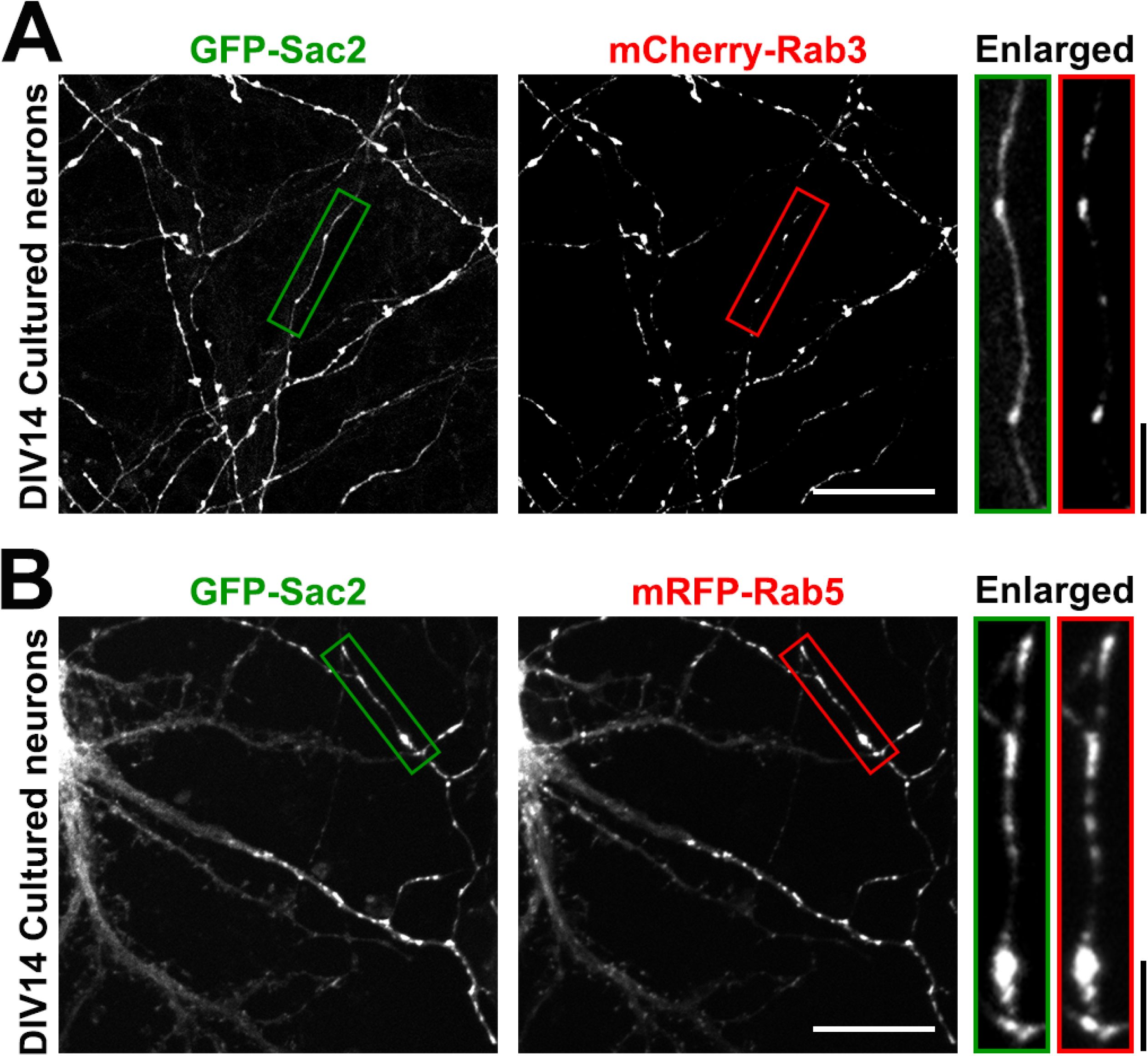
GFP-Sac2 is present in axons and axon terminals. A, B. Cultured WT hippocampal neurons were co-transfected with GFP-Sac2 and either mCherry-Rab3 (marker of axons) (A) or mRFP-Rab5 (B) at DIV8 and imaged at DIV14. The micrographs at right show the areas bracketed by rectangles in the main fields at high magnification. In these enlargements, note the presence of a cytosolic axonal pool of Sac2 in mCherry-Rab3 expressing cells, while in cells expressing of mRFP-Rab5 the cytosolic pool of GFP-Sac2 has relocalized to mRFP-Rab5 positive structures. Scale bar = 10 μm or 5 μm for enlarged images.

### The accumulation of endocytic factors in axon terminals of SJ^RQ^KI neurons is enhanced by the absence of Sac2

We previously reported the occurrence of a very robust exaggerated accumulation of endocytic factors, including clathrin coat components and their endocytic accessory factors, such as amphiphysin and auxilin, in nerve terminals of SJ1^RQ^KI neurons both *in situ* (in all of several brain regions examined) and in neuronal cultures (Cao et al., 2017a). As WT SJ1 does not dephosphorylate PI4P, but its Sac domain does, dephosphorylation of PI(4,5)P_2_ is thought to occur sequentially, with the 5-phosphatase domain acting first. Most likely, this abnormal accumulation of endocytic factors at synapses reflects their impaired shedding from membranes due to the lack of PI4P dephosphorylation by the Sac domain of SJ1.

We performed a similar analysis on neuronal cultures of Sac2KO neurons and Sac2KO/SJ1^RQ^KI double mutant neurons. No such accumulation/clustering was observed in nerve terminals of Sac2KO neurons relative to controls (Fig 3). However, in Sac2KO/SJ1^RQ^KI neurons, accumulation of amphiphysin 2, clathrin light chain (CLC), auxilin and SJ1 itself was strongly more pronounced than in SJ1^RQ^KI neurons, corroborating evidence for a synthetic effect of the two mutations (Fig 3). As in the case of SJ1^RQ^KI neurons (Cao et al., 2017a), the enhanced presynaptic accumulation of endocytic factors observed in double mutant mice was more pronounced at inhibitory than at excitatory presynaptic terminals, which were detected by immunolabeling for vGAT and vGlut1, respectively (Fig EV3).

**Figure 3.**
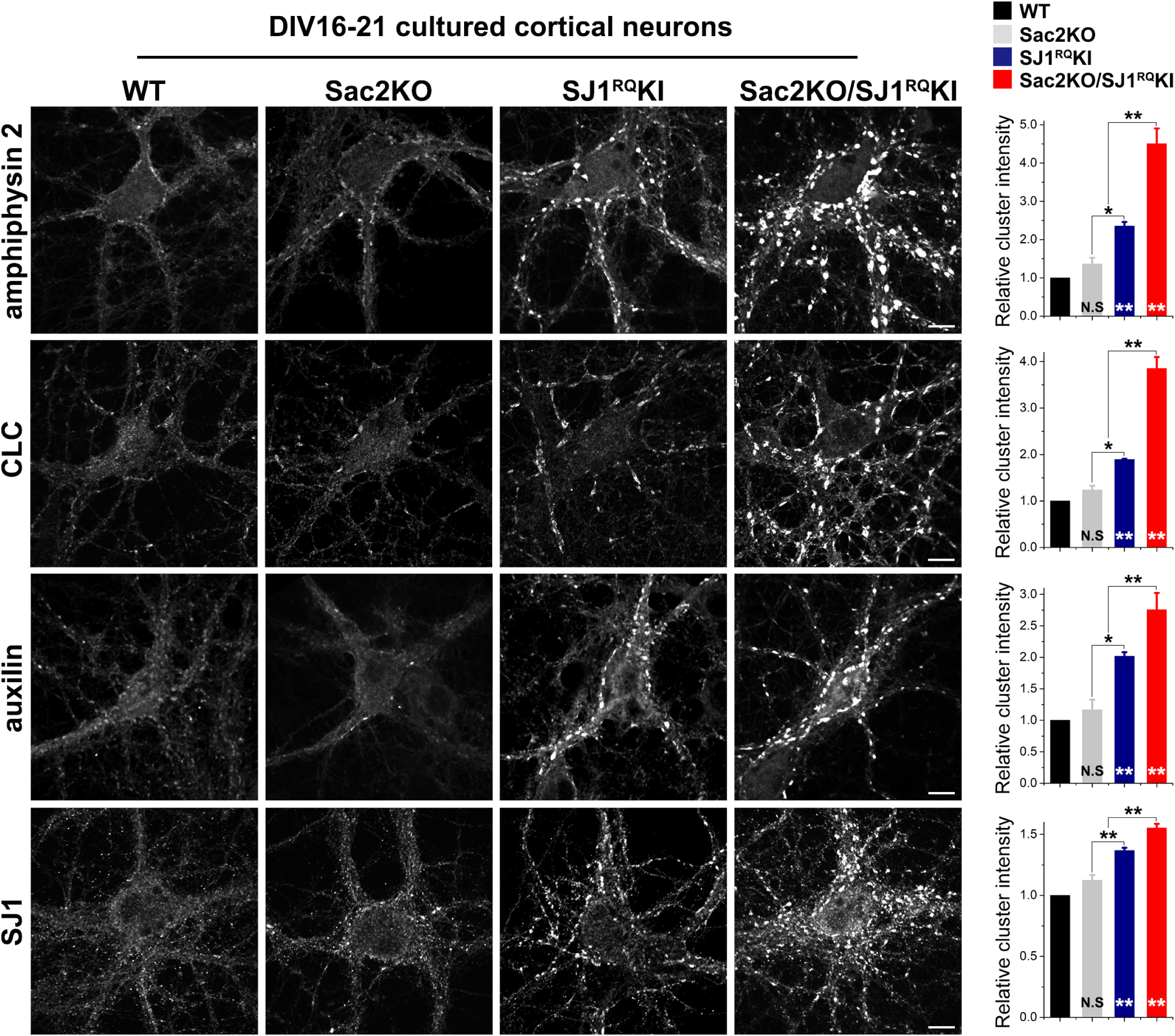
The absence of Sac2 enhances the abnormal accumulation of endocytic factors observed at SJ1^RQ^KI synapses. Left: representative images showing immunoreactivity for amphiphysin 2, clathrin light chain (CLC), auxilin, and SJ1 in DIV16-21 cultured cortical neurons from the genotypes indicated. Scale bar = 10 μm. Right: Quantification of the synaptic clustering of each endocytic protein shown in left panel. amphiphysin 2: WT n = 39, Sac2KO n = 43, SJ1^RQ^KI n = 38 and Sac2KO/SJ1^RQ^KI n = 51 (from 5 independent experiments); CLC: WT n = 36, Sac2KO n = 34, SJ1^RQ^KI n = 32 and Sac2KO/SJ1^RQ^KI n = 34 (from 3 independent experiments); auxilin: WT n = 27, Sac2KO n = 20, SJ1^RQ^KI n = 29 and Sac2KO/SJ1^RQ^KI n = 35 (from 3 independent experiments); SJ1: WT n = 25, Sac2KO n = 22, SJ1^RQ^KI n = 28 and Sac2KO/SJ1^RQ^KI n = 24 (from 3 independent experiments). Data are represented as mean ± SEM; N.S., not significant; *p < 0.05 and **p < 0.01 by ANOVA and Tukey’s post hoc test. Asterisks at the bottom of the bars represent the significance compared to the control.

Since previous studies of Sac2 had revealed that pools of this protein are localized on several endocytic compartments (Hsu et al., 2015, Nakatsu et al., 2015), we also analyzed the distribution of EEA1 (early endosomal marker) and LAMP1 (lysosome marker) in cultured neurons of WT, Sac2KO, SJ1^RQ^KI and Sac2KO/SJ1^RQ^KI mice. However, no obvious difference for any of these markers was observed among the different genotypes and clusters of amphiphysin 2 did not colocalize with these proteins (Fig EV4).

### The accumulation of endocytic factors in axon terminals correlates with the accumulation of abnormal vesicular intermediates

We next performed correlative light and electron microscopy (CLEM) to determine whether the clustered immunoreactivity of endocytic proteins corresponded to the accumulation of endocytic vesicular intermediates. To this aim, SNAP-tagged CLC was transfected into cultured cortical neurons grown on gridded glass coverslips and labeled with the cell permeable Janelia Fluor 549. SNAP-CLC fluorescence colocalized with the prominent clusters of endogenous amphiphysin 2 (Fig EV5A), indicating that this construct is targeted to presynaptic nerve terminals as the endogenous protein. SNAP-CLC positive structures were then marked on the gridded glass coverslips and selected regions were processed for epon embedding, thin sectioning and EM analysis (Fig 4).

**Figure 4.**
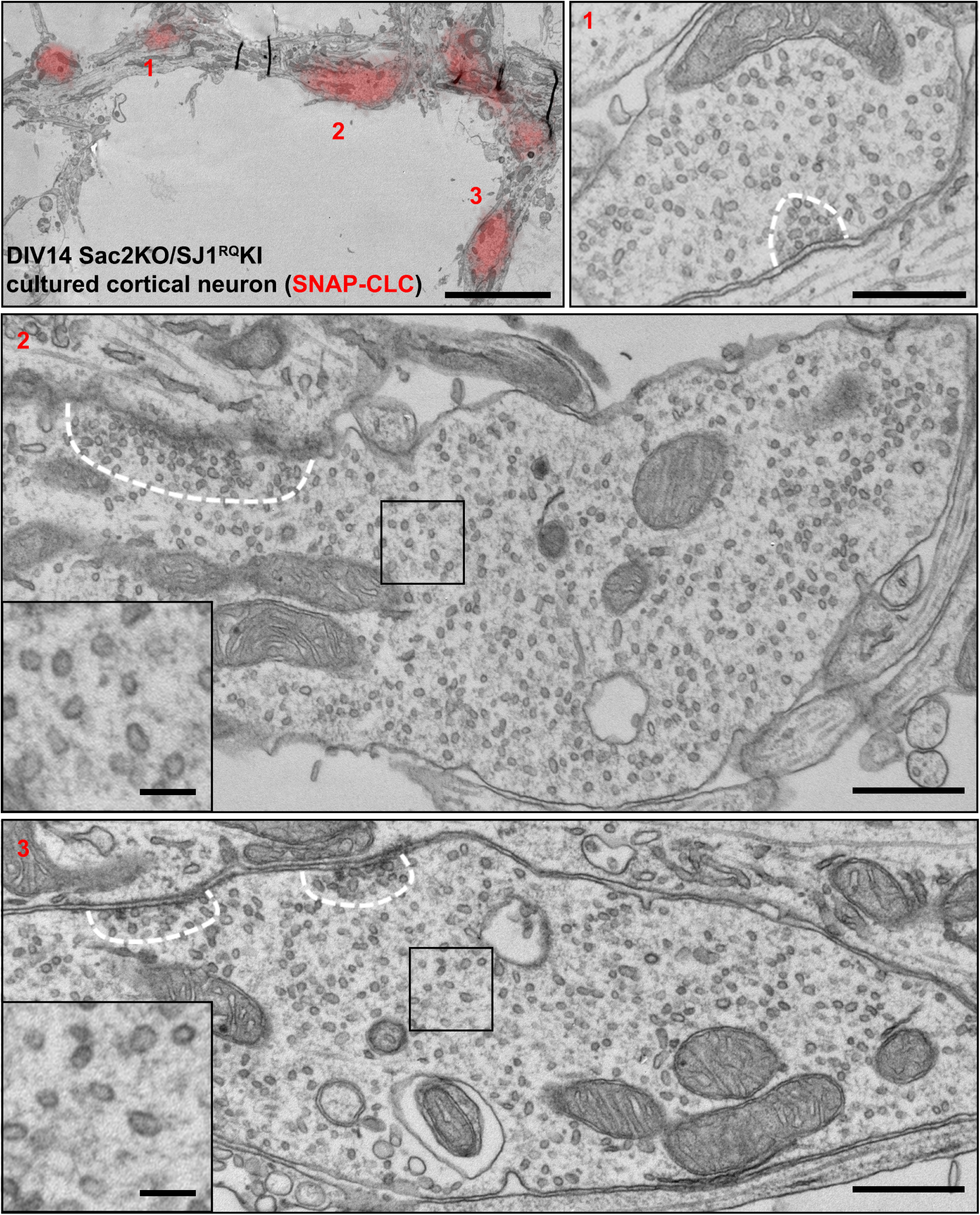
Abnormal accumulation of small vesicles in Sac2KO/SJ1^RQ^KI double mutant presynaptic terminals. Cultured cortical neurons from Sac2KO/SJ1^RQ^KI double mutant mice were transfected with SNAP-CLC and incubated with Janelia Fluor® 549 to perform correlative light and electron microscopy (CLEM). Top left: A low magnification EM image was aligned with the fluorescence image of SNAP-CLC (red). Scale bar = 5 μm. Top right, middle and bottom: High magnification EM images. Note the great abundance of loosely packed small vesicles surrounded by, and intermixed with, a dense protein matrix. A dashed white line outlines the sharp boundary of a classical cluster of tightly packed synaptic vesicles adjacent to a synaptic junction. These vesicles are in the same size range of the surrounding loosely packed vesicles. Scale bar = 500 nm (100 nm for inset).

EM observation revealed that clusters of SNAP-CLC fluorescence in Sac2KO/SJ1^RQ^KI double mutant axon terminals corresponded to large varicose axon terminals as revealed by the massive presence of small vesicles and by the presence of synaptic contacts (Fig 4). However, these accumulations were clearly different from the clusters of bona fide synaptic vesicles of WT synapses (Milovanovic, Wu et al., 2018), because of the larger spacing between them, which was occupied by a dense matrix (Fig 4). In fact, when presynaptic active zones were visible in the section, a sharp boundary was observed between the typically tightly packed synaptic vesicles anchored to these zones (Milovanovic et al., 2018) and the more loosely packed vesicles surrounding them (Fig 4). At lower power EM observation, these accumulations of loosely packed vesicles were reminiscent of the accumulation of clathrin coated vesicles observed by EM around synaptic vesicle clusters at synapses of cultured neurons that either lack SJ1 (SJ1KO neurons) (Cremona et al., 1999, Hayashi et al., 2008) or lack all endophilins (endophilin triple KO neurons) (Milosevic et al., 2011). i.e. the proteins that recruit SJ1 to endocytic sites (Gad, Ringstad et al., 2000, Milosevic et al., 2011). However, in the case of Sac2KO/SJ1^RQ^KI double mutant neurons, clathrin coats could not be clearly observed on the loosely packed vesicles (Fig 4), in spite of evidence for a high concentration of clathrin among them demonstrated by immunofluorescence (Fig 4). Most likely, these vesicles are partially uncoated vesicles, with clathrin and their accessory factors accounting for the matrix surrounding them. Dephosphorylation of PI(4,5)P_2_ at the 5 position, but persistence of PI4P, may be responsible for this phenotype.

Lack of an obvious assembled clathrin coat was also in contrast to what we had observed at synapses of SJ1^RQ^KI adult brains *in situ*, where loosely spaced endocytic vesicular intermediates had a recognizable clathrin coat. However, when we examined nerve terminals of SJ1^RQ^KI neurons in culture, similarly loosely packed vesicles without a clear clathrin coat were observed (Fig EV5B). While the reason for this discrepancy between EM observations at synapses *in situ* and *in vitro* remains to be elucidated, our current findings suggest that the absence of Sac2 simply results in an exaggeration of the trafficking defect of SJ1^RQ^KI, not to a qualitatively different defect.

### Lack of Sac2 accelerates the occurrence of neurodegenerative changes in SJ1^RQ^KI neurons

Previously, we showed that the level of the E3 ubiquitin ligase parkin, another PD gene (PARK2), is robustly increased in SJ1^RQ^KI 6 month-old mouse brains (Cao et al., 2017a). We have now found that this increase is already observed at 3.5 months, but not at 2.5 months (Fig 5A). However, a Sac2KO/SJ1^RQ^KI double mutant mouse that survived for more than two weeks and was sacrificed at 19 days showed a significant increase of parkin in brain relative to brains of similar age from WT, SJ1^RQ^KI or Sac2KO mice (Fig 5B). No increase in parkin was observed in Sac2KO/SJ1^RQ^KI double mouse brain at P0 (Fig 5C). Moreover, lysates from 3-weeks old Sac2KO/SJ1^RQ^KI double mutant neuronal cultures showed upregulation of parkin relative to Sac2KO cultures and to WT and SJ1^RQ^KI cultures of similar age (Fig 5D and E).

**Figure 5.**
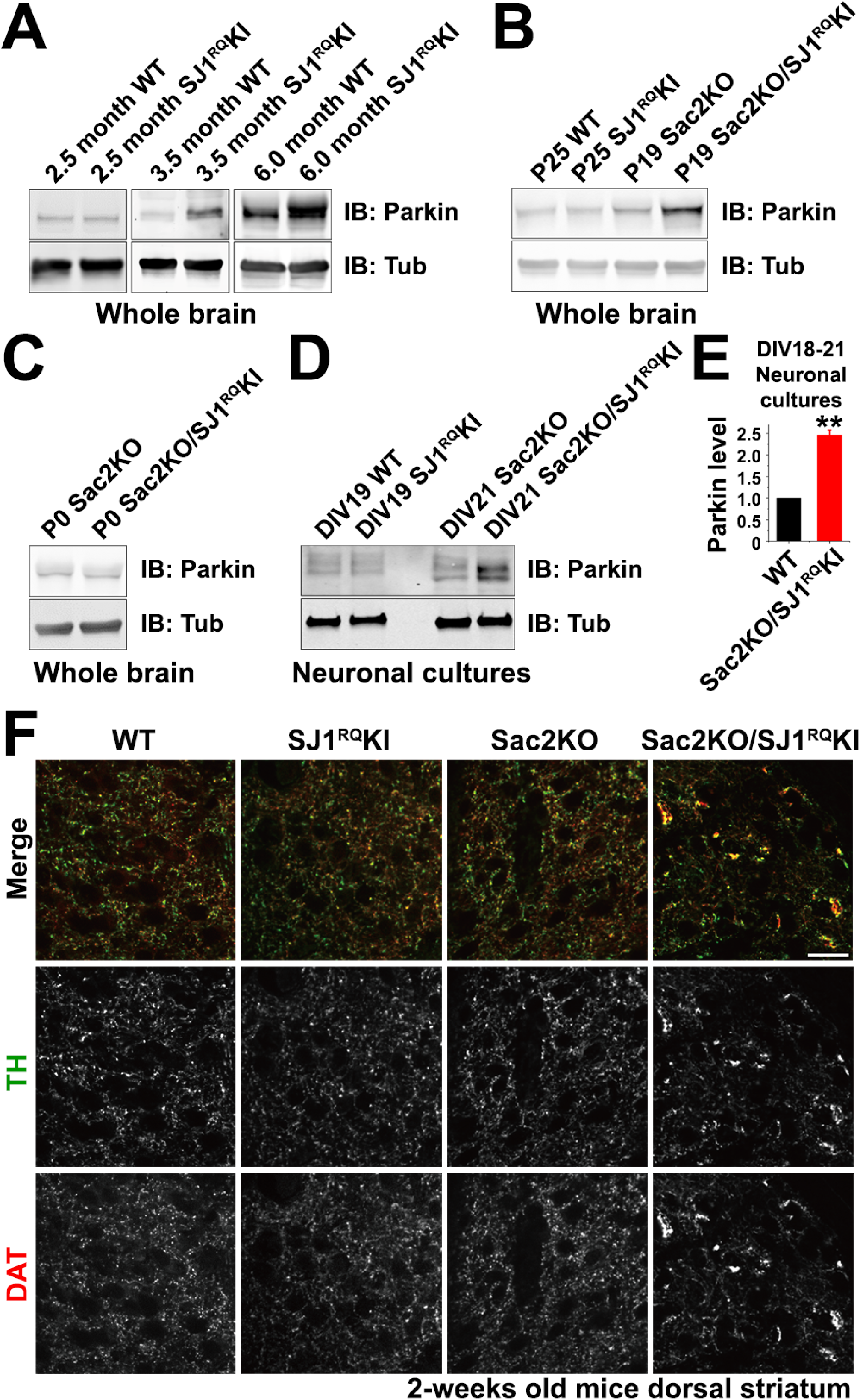
The loss of Sac2 accelerates the occurrence of neurodegenerative phenotypes in SJ1^RQ^KI neurons. A. Level of parkin in SJ1^RQ^KI and a littermate WT control at three different ages as assessed by western blotting. In this field and in the following fields anti-tubulin (Tub) antibodies were used as loading controls. B. Increase of parkin in a P19 (postnatal day 19) Sac2KO/SJ1^RQ^KI double mutant brain relative to brains of a similar age from WT, SJ1^RQ^KI or Sac2KO mice. C. Parkin level was unchanged in P0 Sac2KO/SJ1^RQ^KI brain relative to its littermate control (Sac2KO). D. Representative western blot showing the upregulation of parkin in Sac2KO/SJ1^RQ^KI neuronal cultures relative to Sac2KO and to WT and SJ1^RQ^KI cultures of similar age. E. Quantification of the parkin level in Sac2KO/SJ1^RQ^KI neuronal cultures relative to WT cultures of similar age (DIV18-21). n = 3 (from 2 independent neuronal cultures). Data are represented as mean ± SEM; N.S., not significant; **p < 0.01 by Student’s *t* test. F. Anti-tyrosine hydroxylase (TH, green) and anti-dopamine transporter (DAT, red) immunofluorescence on frozen sections of dorsal striata of 2-week old mice of the genotypes indicated. Aggregation (co-aggregation) of TH and DAT was only observed in Sac2KO/SJ1^RQ^KI double mutant striatum at this age. Scale bar = 20 μm.

This earlier onset of a biochemical parameter that may reflect neurodegeneration was also reflected in an earlier occurrence of structural signs of neurodegeneration. In SJ1^RQ^KI mice, age-dependent dystrophic changes were observed at one month post-natal, but not at 2 weeks, in a subset of axon terminals of nigrostriatal neurons, as revealed by immunofluorescence of frozen sections (Cao et al., 2017a). These changes, which were revealed by abnormal clusters of immunoreactivity for markers of dopaminergic axons, such as tyrosine hydroxylase (TH) or the dopamine plasma membrane transporter (DAT), as well as of a general axonal marker (SNAP25), suggest a special vulnerability of dopaminergic neurons of the nigrostriatal pathway to the SJ1^RQ^ mutation (Cao et al., 2017a). When the same analysis was performed on the rare Sac2KO/SJ1^RQ^KI double mutant mice that survived for two weeks, similar TH- and DAT-positive sparse dystrophic axon terminals were detected in the dorsal striatum of two Sac2KO/SJ1^RQ^KI mice at P15 and P19 days, indicating an earlier occurrence of axonal dystrophy, consistent with a synthetic effect of the absence of Sac2 and of Sac mutation of SJ1 (Fig 5F).

## Discussion

Our results reveal a striking synthetic effect in mice of loss-of-function mutations of two Sac phosphatase domain-containing proteins, SJ1 and Sac2. While mice with the PD inactivating mutation in the Sac domain of SJ1 (SJ^RQ^KI) have neurological defects but generally survive to adulthood and Sac2KO mice do not have an obvious phenotype, the majority of mice with both mutations have severe neurological defects, die perinatally and none survives to adulthood. A synthetic effect is also observed at the cellular level, as absence of Sac2 enhances the presynaptic defects observed in SJ^RQ^KI nerve terminals and anticipates the age at which 1) parkin abnormally accumulates, and 2) dystrophic dopaminergic axons appear in the dorsal striatum. These results suggest that the Sac phosphatase domains of the two proteins have, at least partially, overlapping functions.

Sac2 is an inositol phosphatase preferentially expressed in the brain (Hsu et al., 2015). It was shown to associate with vesicular structures, including Rab5 positive endosomes in non-neuronal cells (Hsu et al., 2015, Nakatsu et al., 2015), and also in proximity of Rab3 positive organelles in pancreatic β-cells (Nguyen, Gandasi et al., 2019), but only little was known about its localization in neurons, although a role in axons had been proposed (Zou et al., 2015). The synergistic effects of the absence of Sac2 on the presynaptic defects produced by the Sac R258Q mutation of SJ1, a protein enriched in nerve terminals, strongly suggest that at least some actions of Sac2 occur at the presynapse. Accordingly, although we could not assess the localization of the endogenous protein, we found that exogenous tagged Sac2 is present in axons and axon terminals and that is recruited to Rab5 positive endosomes.

Nerve terminals of both SJ1^RQ^KI and Sac2KO/SJ1^RQ^KI neuronal cultures are characterized by a striking exaggerated accumulation of clathrin coat components and accessory factors as well as by small vesicles in the size range of synaptic vesicles that surround, but are well separated from, the peculiar clusters of synaptic vesicles (Milovanovic et al., 2018) anchored at presynaptic active zones. These other vesicles are not as tightly packed as typical synaptic vesicle clusters and are separated from each other by a dense matrix. The endocytic factors that accumulate in mutant nerve terminals are likely to be part of the matrix, as strongly suggested by the overlap between CLC fluorescence and these areas revealed by CLEM. Interconversion of phosphoinositide species on synaptic vesicle membranes by phosphorylation-dephosphorylation during their exo-endocytic cycle impacts their traffic by controlling the association-dissociation of cytosolic factors (Cremona & De Camilli, 2001). Thus, these vesicles may represent endocytic intermediates that fail to progress to downstream stations due to the abnormal accumulation of an inappropriate phosphoinositide on their membrane.

The Sac domain of the synaptojanin family can dephosphorylate PI4P, PI3P and PI(3,5)P_2_ *in vitro* (Guo, Stolz et al., 1999), although a main function of this Sac domain in living cells is thought to be the dephosphorylation of PI4P resulting from the action of its 5-phosphatase domain on PI(4,5)P_2_ (Cremona & De Camilli, 2001). The Sac domain of Sac2 acts nearly exclusively of PI4P (Hsu et al., 2015, Nakatsu et al., 2015). Thus, we favor a model in which the combined absence of the two Sac phosphatase activities results in ectopic PI4P accumulation on synaptic membranes. This accumulation, in turn, may prevent complete shedding of endocytic factors whose binding to the membranes is primarily PI(4,5)P_2_ dependent, but may also depend, in part, on PI4P. We note that auxilin, the co-chaperone of HSC70 needed for the disassembly of the clathrin cage after the release of clathrin adaptors, contains a N-terminal PTEN-homology module that binds phospholipids, including PI4P (Guan, Dai et al., 2010). Auxilin is one of the factors abnormally accumulated at SJ^RQ^KI and Sac2KO/SJ1^RQ^KI mutant synapses and may be responsible for the partial uncoating.

Defective endocytic flux along the endocytic pathway due to impaired shedding of endocytic factors may affect reliability of synaptic transmission by resulting in a lower pool of reserve synaptic vesicles. This is what has been observed for mutations in several endocytic factors (Ferguson, Brasnjo et al., 2007, Koh, Verstreken et al., 2004, Luthi, Di Paolo et al., 2001, Milosevic et al., 2011, Zhang, Koh et al., 1998). The occurrence of epileptic seizures in mice with these mutations reflects the greater dependence of inhibitory synapses on synaptic vesicle recycling given their high levels of tonic activity (Evergren, Zotova et al., 2006). Accordingly, even in our present study, inhibitory synapses were the most more severely affected in Sac2KO/SJ1^RQ^KI neurons.

It is also possible that defect in Sac phosphatase activity at synapses may have effects beyond synaptic vesicle recycling. A selective defect in the Sac domain of SJ1 in zebrafish and flies, including the KI of the R258Q mutation in flies, was reported to impact autophagy (George, Hayden et al., 2016, Vanhauwaert, Kuenen et al., 2017). It was proposed that such defect could be explained by abnormal accumulation in nerve terminals of PI3P, a phosphoinositide required for the initial nucleation of the autophagy machinery (Wirth, Joachim et al., 2013). Recent work suggests that PI4P may also be needed at initial stages of autophagy (Judith, Jefferies et al., 2019). Another possibility is that abnormal endocytic traffic may affect availability of ATG9, the transmembrane component of the core autophagy machinery (Ravikumar, Moreau et al., 2010, Webber, Young et al., 2007). The precise role of the Sac domain in autophagy deserves further investigation.

Irrespective of the precise mechanisms through which the combined deficiency of Sac2 and of the Sac domain of SJ1 impact presynaptic function, our results have interesting implications for the PD field. The SJ1 R258Q mutations is responsible for familial (although rare) cases of PD (Krebs et al., 2013, Olgiati et al., 2014, Quadri et al., 2013), and Sac2 has been linked to PD by GWAS studies (Nalls et al., 2014). The functional connection between these two PD-linked proteins revealed by our results points to their shared site of action as a node whose dysfunction may result in PD pathogenesis. Other evidence links SJ1 function to PD. For example, endophilin, which has a major role in recruiting SJ1 to endocytic sites (Gad et al., 2000, Milosevic et al., 2011), binds parkin (Trempe et al., 2009) and loss of endophilin results in elevated levels of parkin (Cao et al., 2014, Trempe et al., 2009). Indeed, both endophilin and SJ1 are ubiquitinated by parkin (Cao et al., 2014, Trempe et al., 2009) and the endophilin A1 gene itself was identified as a risk locus for PD (Chang, Nalls et al., 2017). In addition, parkin is very robustly increased in SJ1^RQ^KI mice brains (Cao et al., 2017a) and here we found that the additional loss of Sac2 accelerates the occurrence of this phenotype. Finally, the PD protein LRRK2 (PARK8) phosphorylates SJ1, endophilin and auxilin (Arranz, Delbroek et al., 2015, Nguyen & Krainc, 2018, Pan, Li et al., 2017). Further elucidating how mutations of these house-keeping proteins impact neuronal functions, and more robustly the function of selective PD-relevant neuronal populations, is expected to advance our understanding of PD mechanisms.

## Materials and Methods

### Animals

SJ1^RQ^KI mice carrying the EOP R258Q mutation were custom generated as reported previously (Cao et al., 2017a). Sac2/INPPF5 KO mice were a gift from Dr. Jonathan Epstein, University of Pennsylvania (Zhu et al., 2009). No obvious neurological defects were observed in Sac2KO mice. Both strains were maintained on the C57BL6/129 genetic background and crossed with each other to generate SJ1/Sac2 double mutant mice. For the comparison of four different genotypes (WT, SJ1^RQ^KI, Sac2KO and SJ1^RQ^KI/Sac2KO), SJ1 heterozygous KI (+/KI) were mated to generate homozygous mice (KI/KI) and WT littermate controls. Sac2KO/SJ1 heterozygous KI (+/KI) double mutant were mated with each other to generate Sac2KO/SJ1^RQ^KI double homozygous and Sac2 single KO littermate controls. All mice were maintained on a 12 h light/dark cycle with standard mouse chow and water ad libitum. All research and animal care procedures were approved by the Yale University Institutional Animal Care and Use Committee.

### Plasmids and Antibodies

The sources of cDNAs were as follows: pEGFPC-Sac2 and pSNAP-human clathrin light chain (CLC) were generated in our lab; pFUGW-GFP-Sac2 was generated from pEGFPC-Sac2 by Janelia Research Campus/HHMI. The following primary antibodies used in this study were generated in our lab: rabbit anti-SJ1, rabbit anti-auxilin. Antibodies obtained from commercial sources were as follows: mouse anti-EEA1 from BD biosciences; mouse anti-amphiphysin 2, rabbit anti-CLC, guinea pig anti-vGluT1 (AB5905), rabbit anti-TH (AB152), and rat anti-DAT (MAB369) from Millipore; rabbit anti-synaptophysin (101 002) and rabbit anti-vGAT (131 002) from Synaptic Systems; rat anti-LAMP1 (ID4) from DSHB; rabbit anti-BAG3 (10599-1-AP) from Proteintech, mouse anti-tubulin (T5168) from Sigma; rabbit anti-tubulin (PA1-21153) from Invitrogen; mouse anti-parkin (4211) from Cell Signaling Technology; Alexa/488-, 594- and 647-conjugated secondary antibodies from Invitrogen. The rabbit anti-Sac2 antibody was a kind gift of Yuxin Mao, (Cornell University, NY).

### Immunoblotting

Post-nuclear supernatant of brain tissue was obtained by homogenization of mouse brain in buffer containing 20 mM Tris, pH 7.4, 150 mM NaCl, and 2 mM EDTA supplemented with protease inhibitors (cOmplete protease Inhibitor cocktail, Roche) and subsequent centrifugation at 700g for 10 mins. Protein concentration was determined by the Pierce BCA Protein Assay Kit. SDS-PAGE and western blotting were performed by standard procedures. Proteins were detected by enhanced chemiluminescence (ECL) reagent and quantified by densitometry using Fiji software.

### Primary neuronal culture and fluorescence microscopy

Cultures of cortical or hippocampal neurons were prepared from P0-P2 neonatal mouse brains as previously described (Ferguson et al., 2007, Park, Lee et al., 2018) and used at DIV14-23. For lenti-virus infection, DIV3 neurons cultured on 12-mm coverslips were infected with 1 μl FUGW-GFP-Sac2 viruses (5E9 Integration Unit (IU)/mL) and fixed after DIV16. Calcium phosphate transfection was performed as previously described (Park et al., 2018). Cells were fixed with 4% formaldehyde [freshly prepared from paraformaldehyde (PFA)] in 0.1 M sodium phosphate pH 7.2 and 4% sucrose, blocked and permeabilized with PBS containing 5% bovine serum albumin (BSA) and 0.1% triton X-100. Primary and secondary antibody incubations for immunofluorescence were subsequently performed in the same buffer. After washing, samples were mounted on slides with Prolong Gold antifade reagent (Invitrogen), observed by either a Perkin Elmer Ultraview spinning disk confocal microscope equipped with 60x CFI PlanApo VC objective or a laser-scanning confocal microscope (LSM 710; Carl Zeiss) equipped with a 63x PlanApo objective.

### Correlative light-electron microscopy (CLEM)

Plasmids encoding SNAP-CLC were electroporated into dissected neuronal suspensions using Amaxa (Lonza) Nucleofector kit at DIV0 before plating on 35mm gridded, glass-bottom MatTek dishes (P35G-1.5-14-CGRD). At DIV14, neurons were stained with 0.5 μM Janelia Fluor® 549 at 37 °C for 1 hr, followed by incubation in original culture medium at 37 °C for 2 hrs before fixation for immunofluorescence or CLEM. Labeled neurons were imaged and their coordinates on the MatTek dishes were recorded using fluorescence microscopy and bright-field DIC, respectively. Then, neurons were fixed with 2.5% glutaraldehyde in 0.1 M sodium cacodylate buffer, post-fixed in 0.1 M sodium cacodylate buffer containing 1% OsO_4_ and 1.5% K_4_Fe(CN)_6_ (Sigma-Aldrich, St. Louis, MO), *en bloc* stained with 2% aqueous uranyl acetate, dehydrated, and embedded in Embed 812. The nerve terminals expressing SNAP-CLC were relocated (based on the pre-recorded coordinates), sectioned and imaged. Ultrathin sections (60-80 nm) were observed in a Philips CM10 microscope at 80 kV, images were taken with iTEM (Soft Imaging System GmbH) and a Morada 1kx1k CCD camera (Olympus). Except when noted, all reagents for EM were from EMS, Hatfield, PA).

### Brain histology

Brain tissues from 2-week old mice were dissected out, immersed immediately in ice cold fixative [4% formaldehyde (see above) in 0.1 M phosphate buffer] and kept in the same fixative overnight at 4 °C. Brains were then transferred to increasing concentrations of sucrose (10%, 20% and 30% (w/v)] in PBS, embedded in OCT (Tissue-Tek) and frozen in liquid nitrogen-cooled isopentane. Coronal or sagittal (15-30 μm thickness) sections were cut with a cryostat and mounted on Superfrost™ Ultra Plus Adhesion slides (Thermo Scientific). Sections were then blocked and permeabilized with a solution containing 3% normal goat serum, 1% BSA, PBS and 0.1% Triton-X100 for 1 hr at room temperature, incubated with primary antibodies (diluted in the same buffer) overnight at 4 °C, washed, incubated with Alexa-conjugated secondary antibodies for 1 hr at room temperature and finally mounted with Prolong Gold antifade reagent with DAPI and sealed with nail polish. Images were acquired with a Perkin Elmer Ultraview spinning disk confocal microscope equipped with 40x objective.

### Quantification of immunoreactivity clustering

Quantification of endocytic protein clustering was performed using Fiji software as follows. After background subtraction, the same threshold intensity was applied to all images and raw intensity values of masked regions were measured by using the “analyze particles” function of Fiji. Quantification of the average fluorescence of puncta bigger than 0.1 μm^2^ was performed for WT control, SJ1^RQ^KI, Sac2KO and Sac2KO/SJ1^RQ^KI neurons.

### Statistical analysis

Unless otherwise specified, data is presented as mean+/-SEM. Statistical significance was determined using the Student’s two-sample t-test for the comparison of two independent groups or ANOVA followed by Tukey’s honest significant difference (HSD) post hoc test for multiple group comparison. p values <0.05, <0.01 are indicated by asterisks * and **, respectively.

## Acknowledgements

We thank Frank Wilson and Rosemary Coolon for outstanding technical assistance, Dr. Jonathan Epstein (University of Pennsylvania) for the gift of Sac2KO mice, Yuxin Mao for the gift of anti-Sac2 antibodies and Kimberley Ritola (Janelia Research Campus) for the generation of lenti-viruses. This work was supported by grants from the NIH (NS36251 and DA018343), the Kavli Foundation, MJFF and the Parkinson’s Foundation to PDC.

## Author contributions

MC and DP performed all the experiments and analysed all the data with the exception of the EM experiments, which were perfomed by YW. MC, DP and PDC wrote the manuscript.

## Expanded View figures

**Fig EV1. Mendelian inheritance of Sac2KO/SJ1^RQ^KI double mutant mice and evidence that Bag3 levels are not affected in the brains of the Sac2 KO mice**

A. The expected normal Mendelian ratio was observed for new-born pups of the genotypes indicated, ruling out embryonic lethality. Number of mice are indicated in parenthesis.

B. Bag3 levels in Sac2KO mice brain. Western blot analysis of whole brain lysates from 4-month-old mice showing that editing of the Sac2 gene did not affect Bag3 expression.

**Fig EV2. Widespread distribution of GFP-Sac2 in all neuronal compartments including presynaptic terminals**

Cultured hippocampal neurons were infected with GFP-Sac2 encoding lentivirus at DIV3 and stained with antibodies to the synaptic vesicle protein synaptophysin (a presynaptic marker) at DIV16. Scale bar = 20 (top), 5 (middle) or 2 (bottom) μm.

**Fig EV3. The clustering of endocytic proteins in Sac2KO/SJ1^RQ^KI double mutant neurons is more severe at GABAergic than at glutamatergic presynaptic nerve terminals**

Triple immunofluorescence staining of DIV16 cultured cortical neurons from Sac2KO/SJ1^RQ^KI mice using for the antigens indicated. Abnormal clusters of amphiphysin 2 overlap more prominently with vGAT positive than with vGLUT1 positive synapses. Scale bar = 10 μm.

**Fig EV4. Normal distribution of early endosome and lysosome markers in Sac2KO/SJ1^RQ^KI double mutant neurons**

Triple immunofluorescence staining of DIV16 cultured cortical neurons from mice of the indicated genotypes. The distribution of EEA1 (green) and LAMP1 (blue) immunoreactivities are normal in the four genotypes, and do not co-localize with the abnormal amphiphysin 2 (red) clusters typical of Sac2KO/SJ1^RQ^KI double mutant neurons. Scale bar = 20 μm.

**Fig EV5. Endocytic intermediates in Sac2KO/SJ1^RQ^KI and SJ1^RQ^KI mice**

A. Colocalization of SNAP-CLC with amphiphysin 2 in nerve terminals of a Sac2KO/SJ1^RQ^KI double mutant neuron. Anti-amphiphysin 2 immunofluorescence (green) of double mutant neurons expressing SNAP-CLC (red). The colocalization of SNAP-CLC with amphiphysin 2 [as observed for endogenous CLC (see Fig 3)] confirms the correct targeting of this construct to synapses. Scale bar = 20 μm.

B. Ultrastructure of a synapse of a cultured SJ1^RQ^KI neuron. EM appearance of a nerve terminal from a cortical neuronal culture (DIV16). As in the case of Sac2KO/SJ1^RQ^KI double mutant cultured neurons, the abundant small vesicles that surround the small remaining synaptic vesicle cluster (white dashed line) are loosely packed and surrounded by a dense matrix, but do not have an obvious clathrin coat. Scale bar = 500 nm (100 nm for inset).

## References

Arranz AM, Delbroek L, Van Kolen K, Guimaraes MR, Mandemakers W, Daneels G, Matta S, Calafate S, Shaban H, Baatsen P, De Bock PJ, Gevaert K, Vanden Berghe P, Verstreken P, De Strooper B, Moechars D (2015) LRRK2 functions in synaptic vesicle endocytosis through a kinase-dependent mechanism. J Cell Sci 128: 541–52

Blauwendraat C, Heilbron K, Vallerga CL, Bandres-Ciga S, von Coelln R, Pihlstrom L, Simon-Sanchez J, Schulte C, Sharma M, Krohn L, Siitonen A, Iwaki H, Leonard H, Noyce AJ, Tan M, Gibbs JR, Hernandez DG, Scholz SW, Jankovic J, Shulman LM et al. (2019) Parkinson’s disease age at onset genome-wide association study: Defining heritability, genetic loci, and alpha-synuclein mechanisms. Mov Disord 34: 866–875

Cao M, Milosevic I, Giovedi S, De Camilli P (2014) Upregulation of Parkin in endophilin mutant mice. J Neurosci 34: 16544–9

Cao M, Wu Y, Ashrafi G, McCartney AJ, Wheeler H, Bushong EA, Boassa D, Ellisman MH, Ryan TA, De Camilli P (2017a) Parkinson Sac Domain Mutation in Synaptojanin 1 Impairs Clathrin Uncoating at Synapses and Triggers Dystrophic Changes in Dopaminergic Axons. Neuron 93: 882–896 e5

Cao YL, Yang YP, Mao CJ, Zhang XQ, Wang CT, Yang J, Lv DJ, Wang F, Hu LF, Liu CF (2017b) A role of BAG3 in regulating SNCA/alpha-synuclein clearance via selective macroautophagy. Neurobiol Aging 60: 104–115

Chang D, Nalls MA, Hallgrimsdottir IB, Hunkapiller J, van der Brug M, Cai F, International Parkinson’s Disease Genomics C, and Me Research T, Kerchner GA, Ayalon G, Bingol B, Sheng M, Hinds D, Behrens TW, Singleton AB, Bhangale TR, Graham RR (2017) A meta-analysis of genome-wide association studies identifies 17 new Parkinson’s disease risk loci. Nat Genet 49: 1511–1516

Cremona O, De Camilli P (2001) Phosphoinositides in membrane traffic at the synapse. J Cell Sci 114: 1041–52

Cremona O, Di Paolo G, Wenk MR, Luthi A, Kim WT, Takei K, Daniell L, Nemoto Y, Shears SB, Flavell RA, McCormick DA, De Camilli P (1999) Essential role of phosphoinositide metabolism in synaptic vesicle recycling. Cell 99: 179–88

Dyment DA, Smith AC, Humphreys P, Schwartzentruber J, Beaulieu CL, Consortium FC, Bulman DE, Majewski J, Woulfe J, Michaud J, Boycott KM (2015) Homozygous nonsense mutation in SYNJ1 associated with intractable epilepsy and tau pathology. Neurobiol Aging 36: 1222 e1–5

Edvardson S, Cinnamon Y, Ta-Shma A, Shaag A, Yim YI, Zenvirt S, Jalas C, Lesage S, Brice A, Taraboulos A, Kaestner KH, Greene LE, Elpeleg O (2012) A deleterious mutation in DNAJC6 encoding the neuronal-specific clathrin-uncoating co-chaperone auxilin, is associated with juvenile parkinsonism. PLoS One 7: e36458

Eisenberg E, Greene LE (2007) Multiple roles of auxilin and hsc70 in clathrin-mediated endocytosis. Traffic 8: 640–6

Evergren E, Zotova E, Brodin L, Shupliakov O (2006) Differential efficiency of the endocytic machinery in tonic and phasic synapses. Neuroscience 141: 123–31

Ferguson SM, Brasnjo G, Hayashi M, Wolfel M, Collesi C, Giovedi S, Raimondi A, Gong LW, Ariel P, Paradise S, O’Toole E, Flavell R, Cremona O, Miesenbock G, Ryan TA, De Camilli P (2007) A selective activity-dependent requirement for dynamin 1 in synaptic vesicle endocytosis. Science 316: 570–4

Fotin A, Cheng Y, Grigorieff N, Walz T, Harrison SC, Kirchhausen T (2004) Structure of an auxilin-bound clathrin coat and its implications for the mechanism of uncoating. Nature 432: 649–53

Gad H, Ringstad N, Low P, Kjaerulff O, Gustafsson J, Wenk M, Di Paolo G, Nemoto Y, Crun J, Ellisman MH, De Camilli P, Shupliakov O, Brodin L (2000) Fission and uncoating of synaptic clathrin-coated vesicles are perturbed by disruption of interactions with the SH3 domain of endophilin. Neuron 27: 301–12

George AA, Hayden S, Stanton GR, Brockerhoff SE (2016) Arf6 and the 5’phosphatase of synaptojanin 1 regulate autophagy in cone photoreceptors. Bioessays 38 Suppl 1: S119–35

Guan R, Dai H, Harrison SC, Kirchhausen T (2010) Structure of the PTEN-like region of auxilin, a detector of clathrin-coated vesicle budding. Structure 18: 1191–8

Guo S, Stolz LE, Lemrow SM, York JD (1999) SAC1-like domains of yeast SAC1, INP52, and INP53 and of human synaptojanin encode polyphosphoinositide phosphatases. J Biol Chem 274: 12990–5

Hardies K, Cai Y, Jardel C, Jansen AC, Cao M, May P, Djemie T, Hachon Le Camus C, Keymolen K, Deconinck T, Bhambhani V, Long C, Sajan SA, Helbig KL, Consortium ARwgotER, Suls A, Balling R, Helbig I, De Jonghe P, Depienne C, et al. (2016) Loss of SYNJ1 dual phosphatase activity leads to early onset refractory seizures and progressive neurological decline. Brain 139: 2420–30

Hayashi M, Raimondi A, O’Toole E, Paradise S, Collesi C, Cremona O, Ferguson SM, De Camilli P (2008) Cell- and stimulus-dependent heterogeneity of synaptic vesicle endocytic recycling mechanisms revealed by studies of dynamin 1-null neurons. Proc Natl Acad Sci U S A 105: 2175–80

Hsu F, Hu F, Mao Y (2015) Spatiotemporal control of phosphatidylinositol 4-phosphate by Sac2 regulates endocytic recycling. J Cell Biol 209: 97–110

Judith D, Jefferies HBJ, Boeing S, Frith D, Snijders AP, Tooze SA (2019) ATG9A shapes the forming autophagosome through Arfaptin 2 and phosphatidylinositol 4-kinase IIIbeta. J Cell Biol 218: 1634–1652

Koh TW, Verstreken P, Bellen HJ (2004) Dap160/intersectin acts as a stabilizing scaffold required for synaptic development and vesicle endocytosis. Neuron 43: 193–205

Koroglu C, Baysal L, Cetinkaya M, Karasoy H, Tolun A (2013) DNAJC6 is responsible for juvenile parkinsonism with phenotypic variability. Parkinsonism Relat Disord 19: 320–4

Krebs CE, Karkheiran S, Powell JC, Cao M, Makarov V, Darvish H, Di Paolo G, Walker RH, Shahidi GA, Buxbaum JD, De Camilli P, Yue Z, Paisan-Ruiz C (2013) The Sac1 domain of SYNJ1 identified mutated in a family with early-onset progressive Parkinsonism with generalized seizures. Hum Mutat 34: 1200–7

Luthi A, Di Paolo G, Cremona O, Daniell L, De Camilli P, McCormick DA (2001) Synaptojanin 1 contributes to maintaining the stability of GABAergic transmission in primary cultures of cortical neurons. J Neurosci 21: 9101–11

Matta S, Van Kolen K, da Cunha R, van den Bogaart G, Mandemakers W, Miskiewicz K, De Bock PJ, Morais VA, Vilain S, Haddad D, Delbroek L, Swerts J, Chavez-Gutierrez L, Esposito G, Daneels G, Karran E, Holt M, Gevaert K, Moechars DW, De Strooper B, et al. (2012) LRRK2 controls an EndoA phosphorylation cycle in synaptic endocytosis. Neuron 75: 1008–21

McPherson PS, Garcia EP, Slepnev VI, David C, Zhang X, Grabs D, Sossin WS, Bauerfeind R, Nemoto Y, De Camilli P (1996) A presynaptic inositol-5-phosphatase. Nature 379: 353–7

Milosevic I, Giovedi S, Lou X, Raimondi A, Collesi C, Shen H, Paradise S, O’Toole E, Ferguson S, Cremona O, De Camilli P (2011) Recruitment of endophilin to clathrin-coated pit necks is required for efficient vesicle uncoating after fission. Neuron 72: 587–601

Milovanovic D, Wu Y, Bian X, De Camilli P (2018) A liquid phase of synapsin and lipid vesicles. Science 361: 604–607

Nakatsu F, Messa M, Nandez R, Czapla H, Zou Y, Strittmatter SM, De Camilli P (2015) Sac2/INPP5F is an inositol 4-phosphatase that functions in the endocytic pathway. J Cell Biol 209: 85–95

Nalls MA, Pankratz N, Lill CM, Do CB, Hernandez DG, Saad M, DeStefano AL, Kara E, Bras J, Sharma M, Schulte C, Keller MF, Arepalli S, Letson C, Edsall C, Stefansson H, Liu X, Pliner H, Lee JH, Cheng R et al. (2014) Large-scale meta-analysis of genome-wide association data identifies six new risk loci for Parkinson’s disease. Nat Genet 46: 989–93

Nguyen M, Krainc D (2018) LRRK2 phosphorylation of auxilin mediates synaptic defects in dopaminergic neurons from patients with Parkinson’s disease. Proc Natl Acad Sci U S A 115: 5576–5581

Nguyen PM, Gandasi NR, Xie B, Sugahara S, Xu Y, Idevall-Hagren O (2019) The PI(4)P phosphatase Sac2 controls insulin granule docking and release. J Cell Biol 218: 3714–3729

Olgiati S, De Rosa A, Quadri M, Criscuolo C, Breedveld GJ, Picillo M, Pappata S, Quarantelli M, Barone P, De Michele G, Bonifati V (2014) PARK20 caused by SYNJ1 homozygous Arg258Gln mutation in a new Italian family. Neurogenetics 15: 183–8

Olgiati S, Quadri M, Fang M, Rood JP, Saute JA, Fen Chien H, Bouwkamp CG, Graafland J, Minneboo M, Breedveld GJ, Zhang J, International Parkinsonism Genetics N, Verheijen FW, Boon AJ, Kievit AJ, Jardim LB, Mandemakers W, Barbosa ER, Rieder CR, Leenders KL, et al. (2015) DNAJC6 mutations associated with early-onset Parkinson’s disease. Ann Neurol

Pan PY, Li X, Wang J, Powell J, Wang Q, Zhang Y, Chen Z, Wicinski B, Hof P, Ryan TA, Yue Z (2017) Parkinson’s Disease-Associated LRRK2 Hyperactive Kinase Mutant Disrupts Synaptic Vesicle Trafficking in Ventral Midbrain Neurons. J Neurosci 37: 11366–11376

Park D, Lee U, Cho E, Zhao H, Kim JA, Lee BJ, Regan P, Ho WK, Cho K, Chang S (2018) Impairment of Release Site Clearance within the Active Zone by Reduced SCAMP5 Expression Causes Short-Term Depression of Synaptic Release. Cell Rep 22: 3339–3350

Quadri M, Fang M, Picillo M, Olgiati S, Breedveld GJ, Graafland J, Wu B, Xu F, Erro R, Amboni M, Pappata S, Quarantelli M, Annesi G, Quattrone A, Chien HF, Barbosa ER, International Parkinsonism Genetics N, Oostra BA, Barone P, Wang J et al. (2013) Mutation in the SYNJ1 gene associated with autosomal recessive, early-onset Parkinsonism. Hum Mutat 34: 1208–15

Ravikumar B, Moreau K, Jahreiss L, Puri C, Rubinsztein DC (2010) Plasma membrane contributes to the formation of pre-autophagosomal structures. Nat Cell Biol 12: 747–57

Schuske KR, Richmond JE, Matthies DS, Davis WS, Runz S, Rube DA, van der Bliek AM, Jorgensen EM (2003) Endophilin is required for synaptic vesicle endocytosis by localizing synaptojanin. Neuron 40: 749–62

Trempe JF, Chen CX, Grenier K, Camacho EM, Kozlov G, McPherson PS, Gehring K, Fon EA (2009) SH3 domains from a subset of BAR proteins define a Ubl-binding domain and implicate parkin in synaptic ubiquitination. Mol Cell 36: 1034–47

Vanhauwaert R, Kuenen S, Masius R, Bademosi A, Manetsberger J, Schoovaerts N, Bounti L, Gontcharenko S, Swerts J, Vilain S, Picillo M, Barone P, Munshi ST, de Vrij FM, Kushner SA, Gounko NV, Mandemakers W, Bonifati V, Meunier FA, Soukup SF et al. (2017) The SAC1 domain in synaptojanin is required for autophagosome maturation at presynaptic terminals. EMBO J 36: 1392–1411

Verstreken P, Koh TW, Schulze KL, Zhai RG, Hiesinger PR, Zhou Y, Mehta SQ, Cao Y, Roos J, Bellen HJ (2003) Synaptojanin is recruited by endophilin to promote synaptic vesicle uncoating. Neuron 40: 733–48

Webber JL, Young AR, Tooze SA (2007) Atg9 trafficking in Mammalian cells. Autophagy 3: 54–6

Wirth M, Joachim J, Tooze SA (2013) Autophagosome formation--the role of ULK1 and Beclin1-PI3KC3 complexes in setting the stage. Semin Cancer Biol 23: 301–9

Zhang B, Koh YH, Beckstead RB, Budnik V, Ganetzky B, Bellen HJ (1998) Synaptic vesicle size and number are regulated by a clathrin adaptor protein required for endocytosis. Neuron 21: 1465–75

Zhu W, Trivedi CM, Zhou D, Yuan L, Lu MM, Epstein JA (2009) Inpp5f is a polyphosphoinositide phosphatase that regulates cardiac hypertrophic responsiveness. Circ Res 105: 1240–7

Zou Y, Stagi M, Wang X, Yigitkanli K, Siegel CS, Nakatsu F, Cafferty WB, Strittmatter SM (2015) Gene-Silencing Screen for Mammalian Axon Regeneration Identifies Inpp5f (Sac2) as an Endogenous Suppressor of Repair after Spinal Cord Injury. J Neurosci 35: 10429–39

